# Winter nutrients constrain spring drawdown, but only weakly predict bloom intensity in the western Irish Sea

**DOI:** 10.64898/2026.10.27.762229

**Authors:** William Ross Hunter, Eugene O’Kane, Claire Smyth, Daniel McClements, Matthew Service, Adam Mellor

## Abstract

Phytoplankton blooms link climatic conditions to marine food webs and carbon cycling, yet relationships between climate forcing and bloom magnitude remain weak and inconsistent in shelf seas. Using a multi-decadal, high-frequency observational record from the western Irish Sea gyre, we investigated how climate variability influences nutrient dynamics and spring phytoplankton bloom development. Sea surface temperature, marine heatwave intensity and the North Atlantic Oscillation exhibited weak and inconsistent relationships with bloom intensity. However, climatic influences were more evident in nutrient dynamics, particularly the timing of seasonal nutrient drawdown, whereas the magnitude of nutrient utilisation was strongly constrained by winter nutrient inventories. Translation of nutrient drawdown into phytoplankton biomass showed marked attenuation: nitrate consumption explained little variability in bloom intensity, whereas silicate drawdown exhibited a moderate, non-deterministic association consistent with functional constraints on diatom growth and community structure. Our results reveal relationships between climate variability and ecosystem function weaken across successive ecosystem levels. While nutrient inventories provide strong constraints on nutrient use, additional ecological and stoichiometric processes influence how nutrient uptake is translated into phytoplankton biomass. Our findings suggest that weak climate-bloom relationships may arise from hierarchical ecological filtering and highlight the importance of biological constraints when assessing ecosystem responses to climate variability in shelf seas.

## Introduction

Phytoplankton blooms are a defining feature of productive marine ecosystems and play a central role in linking climate variability to ocean biogeochemistry, food web dynamics, and carbon cycling (Falkowski et al. 1998, Arrigo 2005). In temperate shelf seas, the spring phytoplankton bloom accounts for between 30 and 60 % of annual primary production during a brief seasonal window (Behrenfeld et al. 2006, Sharples et al. 2006, Platt et al. 2008). As such, considerable attention has been directed toward understanding how climate change and anthropogenic nutrient inputs influence bloom phenology, magnitude, and composition (Edwards and Richardson 2004, Friedland et al. 2024). Over past decades, there has been considerable concern about nutrient enrichment in European shelf seas, driven by terrestrial runoff, riverine inputs, and atmospheric deposition associated with agricultural and industrial activity (Lotze et al. 2006, Halpern et al. 2008, Ural-Janssen et al. 2024). Elevated concentrations of dissolved nitrogen (N), phosphorus (P), and silica (Si) can promote phytoplankton growth and alter community structure (Tilman 1977, Egge and Aksnes 1992, Makareviciute-Fichtner et al. 2024), motivating large scale monitoring programmes and regulatory frameworks such as the EU Water Framework Directive and Marine Strategy Framework Directive (European and Council 2000, 2008, Ferreira et al. 2011). However, despite sustained investment in monitoring and management, relationships between nutrient enrichment and phytoplankton blooms have often proven inconsistent, particularly when assessed across interannual timescales (Cloern and Jassby 2008, Planque 2016).

Phytoplankton blooms are not controlled solely by nutrient availability (Sverdrup 1953, Behrenfeld 2010). Instead, their dynamics are governed by the combined effects of physical forcing, nutrient cycling, physiological constraints on nutrient use, and ecological processes such as grazing and community composition (Cullen 1991). Climatic variability can regulate the timing of stratification, governing nutrient access through its influence on surface forcing and mixing (Sverdrup 1953, Sharples et al. 2006, Behrenfeld 2010). Winter nutrient availability sets the upper limit on potential seasonal primary production; although the extent to which these nutrients are subsequently converted into phytoplankton biomass may depend upon the physical and ecological processes operating during bloom development (Sharples et al. 2006). The conversion of nutrient uptake into phytoplankton biomass is mediated by stoichiometric requirements and ecosystem structure, such that nutrient drawdown may not scale linearly with changes in phytoplankton biomass (Droop 1973, Arrigo 2005, Behrenfeld 2010). Consequently, climate signals may be attenuated through the ecosystem, reducing predictability of phytoplankton bloom expression even when the physical and biogeochemical controls are well defined (Behrenfeld and Boss 2014, Payne et al. 2016, Planque 2016).

Shelf seas account for a disproportionate share of global marine primary production (Bauer et al. 2013), are subject to strong anthropogenic pressures (Lotze et al. 2006, Halpern et al. 2008), and exhibit pronounced spatial gradients in physical structure and nutrient supply (Hill et al. 1997, Horsburgh et al. 2000). These characteristics create strong spatiotemporal variability in ecosystem functioning, and relationships among climate forcing, nutrient availability and phytoplankton bloom expression are often weak, variable and difficult to predict (Cloern and Jassby 2008, Payne et al. 2016). Consequently, shelf sea systems provide a natural laboratory for investigating how climate signals are propagated, transformed and potentially attenuated across successive stages of ecosystem functioning. The Irish Sea is a semi enclosed temperate shelf sea bounded by Great Britain and Ireland, connected to the adjoining Atlantic shelf through the North Channel and St George’s Channel (Knight and Howarth 1999, Xing and Davies 2001, Fernand et al. 2006). While much of the basin is shallow and tidally mixed, the central and western Irish Sea reaches depths of approximately 120 m and experiences weaker tidal mixing (Hill et al. 1997, Horsburgh et al. 2000). During spring and summer, surface heating leads to thermal stratification, isolating a dome of cooler bottom water and establishing a cyclonic gyre with limited lateral exchange (Horsburgh et al. 2000). The western Irish Sea gyre experiences recurrent seasonal nutrient depletion, pronounced spring phytoplankton blooms and substantial interannual climatic variability, making it an ideal system for investigating the coupling between climate forcing, nutrient dynamics and phytoplankton bloom development (Dickey-Collas et al. 1996, Hill et al. 1997, Gowen et al. 1998, Jardine et al. 2022).

Since the 1990s, the Agri Food and Biosciences Institute (AFBI) has maintained a long term oceanographic observatory within the western Irish Sea gyre, providing high frequency measurements of dissolved inorganic nutrients and associated physical variables (Gowen et al. 2002). The observatory includes both a long-term moored monitoring programme, which has generated continuous records of dissolved inorganic nutrients and temperature since 1997, and a ship-based CTD-rosette sampling programme that provides chlorophyll and discrete water-quality measurements. A consistent and quality-controlled archive of CTD-rosette observations is available from 2006 onwards. Together, these complementary observations offer a valuable opportunity to investigate how climate variability, nutrient supply and biological processes interact to shape phytoplankton bloom dynamics over timescales relevant to ecosystem functioning and management (Payne et al. 2016, Planque 2016). Here we investigate the coupling between climate forcing, nutrient cycling and the spring phytoplankton bloom in a temperate shelf-sea system, using long-term nutrient and temperature observations spanning 1997-2024 together with chlorophyll and associated biological measurements available for 2006-2024. We test how: (i) climate variability influences the timing of seasonal nutrient drawdown; (ii) the magnitude of seasonal nutrient drawdown is determined by winter nutrient availability; and (iii) spring bloom intensity is regulated more strongly by biological processes than by nutrient drawdown alone. We hypothesise that climate signals are progressively attenuated as they propagate through physical, chemical and biological processes, resulting in a weakening of relationships from climate, to nutrient dynamics to phytoplankton bloom expression. Our study does not seek to improve empirical prediction of bloom magnitude. Instead, it focuses on understanding why such prediction is often unsuccessful in temperate shelf seas (Payne et al. 2016, Planque 2016). Through this approach we seek to provide a mechanistic explanation for the weak and variable relationships often observed between climate forcing and phytoplankton bloom expression in shelf sea ecosystems (Behrenfeld and Boss 2014, Friedland et al. 2024).

## Methods

### Data sources and nutrient time-series reconstruction

We analysed long-term physical, chemical, and biological observations collected at the Station 38A oceanographic observatory in the western Irish Sea (53° 46.830′ N, 05° 38.030′ W). The observational record comprised high-frequency moored measurements of dissolved inorganic nutrients and temperature collected between 1997 and 2023, supplemented by ship-based CTD-rosette observations collected between 2006 and 2024. Chlorophyll and phytoplankton bloom metrics were derived from the CTD-rosette sampling programme, and therefore analyses involving bloom phenology and bloom intensity were restricted to the period 2006-2024. Station 38A is located within a seasonally stratified gyre at a nominal water depth of ∼94 m over fine muddy sediments.

For over three decades, the Agri-Food and Biosciences Institute (AFBI) maintained an instrumented subsurface mooring at Station 38A. The mooring comprised a McLane® Remote Access Sampler (RAS-100, McLane Research Laboratories, Falmouth, MA, USA) and a Sea-Bird SBE19 conductivity-temperature-depth (CTD) recorder deployed at ∼20 m depth. The system collected daily water samples preserved with mercuric chloride for dissolved inorganic nutrient analysis (nitrate, nitrite, ammonium, phosphate, and silicate). Ship-based CTD-rosette sampling was conducted throughout much of the observatory’s history; however, a consistent and quality-controlled archive of chlorophyll and discrete water-sample observations was only available from 2006 onwards. Analyses requiring these measurements were therefore restricted to the 2006-2024 period.

The mooring was serviced approximately every six weeks by research cruises aboard RV Corystes, during which stored samples were recovered and the system inspected. Ship based CTD casts were conducted during each visit using a Sea Bird SBE9 CTD and rosette sampler, with discrete water samples collected near the seabed, at intermediate depth (∼40 m), and near the surface for laboratory nutrient analysis. These observations provided depth resolved context for the moored measurements. High frequency nutrient data from the RAS 100 sampler (1997–2023) were treated as primary observations. Using this record, a daily climatology was constructed independently for each nutrient by calculating mean concentrations for each day of year across the full sampling period, capturing their seasonal structure while smoothing short term episodic variability.

The mooring was removed in February 2023 due to persistent limitations in site access. For the post mooring period (2023–2025), depth averaged (0–40 m) nutrient concentrations were collected every six weeks using a Seabird® 911 CTD and Rosette water sampler. These data were used to constrain climatological values. Systematic differences between fixed depth moored observations and depth integrated CTD measurements were accounted for by bias correcting CTD concentrations using linear regressions fitted during periods of temporal overlap. Bias corrected CTD values replaced climatological estimates on sampling days; climatological values were retained on unsampled days. Where multiple observations occurred on a single day, values were averaged, and data provenance was tracked explicitly.

This approach yields a continuous, gap free daily nutrient time series that preserves the long term seasonal structure observed in the historical moored record while incorporating direct observations where available. Use of a site specific climatology derived from high frequency in situ measurements ensures that reconstructed values remain anchored to local hydrographic and biogeochemical conditions rather than relying on regional climatology or model outputs that may not resolve site scale dynamics. To evaluate the robustness of this approach, we conducted sensitivity analyses examining the influence of climatological gap-filling, bias correction, and sampling frequency on derived drawdown and bloom metrics; these analyses are presented in the Supplementary Methods.

The reconstructed time series does not resolve short term episodic variability associated with storms, transient mixing, or short lived biological events; post 2023 values on unsampled days therefore represent background seasonal conditions rather than exact daily states. Bias correction further assumes a stable relationship between depth integrated and fixed depth measurements through time. Despite these limitations, the dataset provides a robust foundation for analyses at seasonal to interannual timescales, which are the focus of this study.

### Nutrient sampling and laboratory analysis

Nutrient samples were collected either using a McLane® RAS 100 automated water sampler deployed on the mooring or from discrete Niskin bottles mounted on a CTD rosette during ship based surveys. Upon recovery of the RAS sampler, preserved water samples were removed from storage bags and subsampled, with 20 mL passed through sterile 0.22 µm filters to remove particulate material before being transferred into sterile, single use 25 mL polypropylene containers (ThermoFisher Scientific Sterilin® Certified Universal Sample Containers). Water samples collected from the rosette water sampler were treated using the same procedure, with filtration through sterile 0.22 µm filters prior to transfer into identical Sterilin® containers. All samples were stored frozen at −20 °C and transported frozen to the laboratory. Prior to analysis, samples were thawed at room temperature over approximately three hours.

All nutrient analyses were conducted at the AFBI Marine Chemistry Laboratory using a SEAL Analytical QuAAtro segmented flow analyser, following established colorimetric methods for dissolved inorganic nutrients (Cheeseman and Wilson 1978). Dissolved ammonium was determined using the Berthelot reaction, in which ammonia reacts with hypochlorite under alkaline conditions to form monochloramine, followed by reaction with phenol in the presence of nitroprusside ions to produce indophenol blue, quantified colorimetrically at 630 nm. Total oxidised nitrogen (nitrate + nitrite) was quantified after reduction of nitrate to nitrite using a copper–cadmium reduction column, followed by diazotisation with sulphanilamide and coupling with N(1 naphthyl)ethylenediamine dihydrochloride to form an azo dye measured at 550 nm. Dissolved nitrate concentrations were calculated as the total oxidised nitrogen concentration minus measured nitrite. Dissolved phosphate and silicate were determined using analogous standardised colorimetric procedures. All nutrient concentrations are reported in micromoles per litre (µmol L⁻¹).

The AFBI Marine Chemistry Laboratory operates under ISO 9001 accreditation and participates in the QUASIMEME inter laboratory quality assurance and control scheme, ensuring analytical traceability and comparability with wider marine biogeochemical datasets. Analytical precision and accuracy were maintained through routine calibration, use of certified reference materials, and internal laboratory standards in accordance with established QA/QC protocols. All nutrient data used in subsequent analyses therefore derive from validated measurements produced under accredited analytical procedures.

### Temperature data and derivation of climate metrics

Near-surface seawater temperature data were obtained from daily mean measurements recorded by a Sea-Bird Scientific SBE56 temperature sensor deployed at approximately 2 m water depth on the S38A oceanographic mooring. These data are publicly available via the ICES Report on Ocean Climate data portal (https://iroc.ices.dk/). Raw temperature observations were aggregated to daily means prior to analysis, yielding a continuous daily time series of near-surface thermal conditions representative of the mixed-layer environment experienced by phytoplankton during bloom development. All temperature-regime descriptors used in this study were derived from this daily temperature measurements.

Marine heatwave and cold spell metrics were calculated following the threshold based framework of Hobday et al. (2016), which defines thermal extremes relative to a seasonally varying climatological baseline. Analyses were implemented using the RMarineHeatwaves package (Schlegel and Smit 2018), which identifies discrete heatwave and cold spell events and quantifies their duration and intensity. Cumulative annual intensity metrics were derived by integrating anomaly magnitudes across all detected events within each year, providing a measure of sustained thermal forcing rather than short lived extremes.

Mean seasonal sea surface temperature was calculated over predefined winter and spring periods corresponding to nutrient reset and bloom development phases, respectively. Large-scale atmospheric forcing was characterised independently using the winter-mean North Atlantic Oscillation (NAO) index averaged over December–March (DJFM), which was treated as a contextual indicator of regional climate state rather than a direct driver of biological responses. Temperature-derived metrics were evaluated primarily in relation to nutrient drawdown timing and, diagnostically, for potential indirect associations with biological structure.

### Data Treatment

Analyses of bloom phenology and bloom intensity were restricted to 2006-2024, corresponding to the period for which a consistent and quality-controlled archive of chlorophyll observations was available from the CTD-rosette monitoring programme. Nutrient and temperature observations from the moored record (1997-2023) were used to characterise long-term seasonal and interannual variability and to construct nutrient climatologies. Bloom phenology and bloom intensity metrics were derived from chlorophyll observations collected during the CTD-rosette monitoring programme (2006-2024).

#### Definition of seasonal periods

Winter conditions were defined as the late winter nutrient-replete period (January-February), when nutrient concentrations are typically replenished through seasonal vertical mixing and biological uptake is minimal. Spring conditions were defined as the period of seasonal phytoplankton development (April-June), during which nutrient drawdown and biomass accumulation occur. Nutrient drawdown phenology was assessed during the transitional spring period (March-June), capturing the progression from winter nutrient repletion to active biological utilisation.

#### Nutrient drawdown metrics

For each nutrient, seasonal utilisation was characterised using two complementary metrics. Drawdown timing was defined as the first day of year on which concentrations declined to 75% of the January-February mean concentration. Drawdown magnitude was calculated as the difference between the January-February mean concentration and the subsequent minimum concentration observed during March-June. These metrics respectively quantify the timing and extent of seasonal nutrient depletion.

#### Bloom intensity metrics and biological variables

Spring bloom intensity was quantified as the difference between the winter minimum chlorophyll *a* concentration (January-February) and the spring maximum chlorophyll *a* concentration (April-June), providing an integrated measure of seasonal biomass accumulation. Bloom metrics were calculated annually for the period 2006-2024. Relationships between nutrient drawdown and bloom intensity were examined to assess how seasonal nutrient utilisation translated into phytoplankton biomass development. Nitrate drawdown was evaluated as an indicator of seasonal nitrogen utilisation, while silicate drawdown was examined separately because of the obligate silicate requirement of diatoms and their importance within temperate spring blooms. Biological structure was characterised using annual maximum diatom abundance, which was used to aid interpretation of variations in bloom intensity and nutrient use.

#### Climate–biology associations

Relationships between climate metrics (mean sea surface temperature, cumulative marine heatwave intensity, and the winter North Atlantic Oscillation index) and maximum diatom abundance were examined to assess potential indirect influences of climate variability on ecosystem composition. These analyses were intended to support conceptual interpretation of ecological pathways linking climate forcing, nutrient dynamics, and bloom development, rather than to infer direct causal mechanisms.

### Statistical analyses

All analyses and figure production were performed using reproducible scripts in R (R CoreTeam 2024). Data ingestion and processing used the packages readr and dplyr (Wickham et al. 2024), and tidyr (Wickham et al. 2024). Relationships between climate variables, nutrient metrics, and biological responses were evaluated using linear models, consistent with the aim of quantifying first order associations and maintaining comparability across nutrients and processes. Linear models were applied uniformly to characterise relationships between climate forcing and nutrient drawdown timing, winter nutrient availability and drawdown magnitude, and nutrient drawdown and bloom intensity.

Generalised additive models (GAMs) were additionally fitted as an exploratory diagnostic to assess whether linear models masked substantial nonlinear structure in relationships where nonlinear biological responses might plausibly be expected. GAMs were implemented using the mgcv package (Wood 2017) with restricted smoothers (basis dimension k = 4) and REML estimation to limit model flexibility given the small sample sizes and to avoid over fitting. GAMs were used solely for diagnostic comparison with linear models and were not employed as the primary basis for inference (Supplementary Table 2).

## Results

Substantial interannual variability was observed across physical forcing, nutrient concentrations, and spring bloom intensity over the study period (Figure 1). Winter thermal conditions and climate indices varied markedly among years, as did winter nutrient concentrations and the magnitude of spring biomass accumulation. However, bloom intensity did not show a simple or consistent correspondence with interannual variability in climate metrics alone, suggesting that bloom expression is mediated by intermediate processes rather than responding directly to climatic forcing.

**Figure 1.**
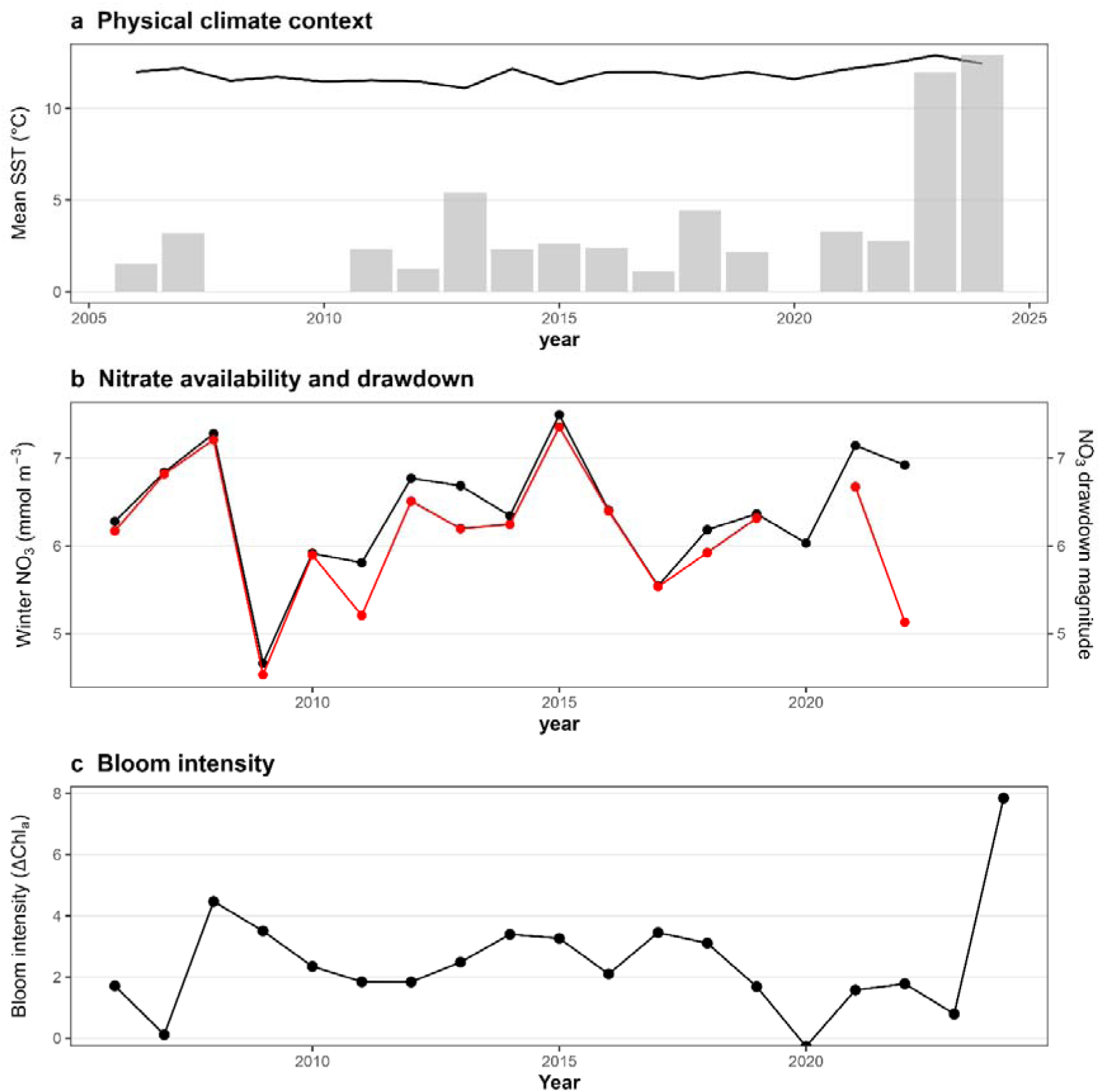
Interannual variability in physical forcing, nutrient dynamics, and spring bloom intensity. **a,** mean sea surface temperature (line) and cumulative heatwave intensity (bars) illustrate interannual variability in the physical environment. **b,** Winter nitrate concentration (black) and subsequent seasonal nitrate drawdown magnitude (red) show year to year variability in nutrient availability and depletion. **c,** Spring bloom intensity, expressed as the difference between winter minimum and spring maximum chlorophyll, varies strongly among years.

The onset of seasonal nutrient drawdown was characterised using separate metrics of drawdown timing and drawdown magnitude, allowing variation in these components of nutrient cycling to be evaluated independently (Figure 2). Drawdown timing varied among years and nutrients but showed only weak associations with mean seasonal sea surface temperature. Across nutrients, linear relationships between drawdown timing and mean SST were weak (R² < 0.05 in all cases; Supplementary Table 3), indicating that interannual variation in the seasonal onset of nutrient utilisation was not tightly prescribed by near-surface thermal conditions alone. In contrast, drawdown magnitude was strongly related to winter nutrient availability. Across nutrients, linear models demonstrated positive relationships between winter mean concentration and subsequent drawdown magnitude, with explanatory power ranging from moderate to very high (R² ≈ 0.31–0.91, p < 0.05; Supplementary Table 3). These results indicate that while climate variability may influence the timing of nutrient drawdown, the total quantity of nutrients consumed during the spring bloom is more closely related to winter nutrient supply.

**Figure 2.**
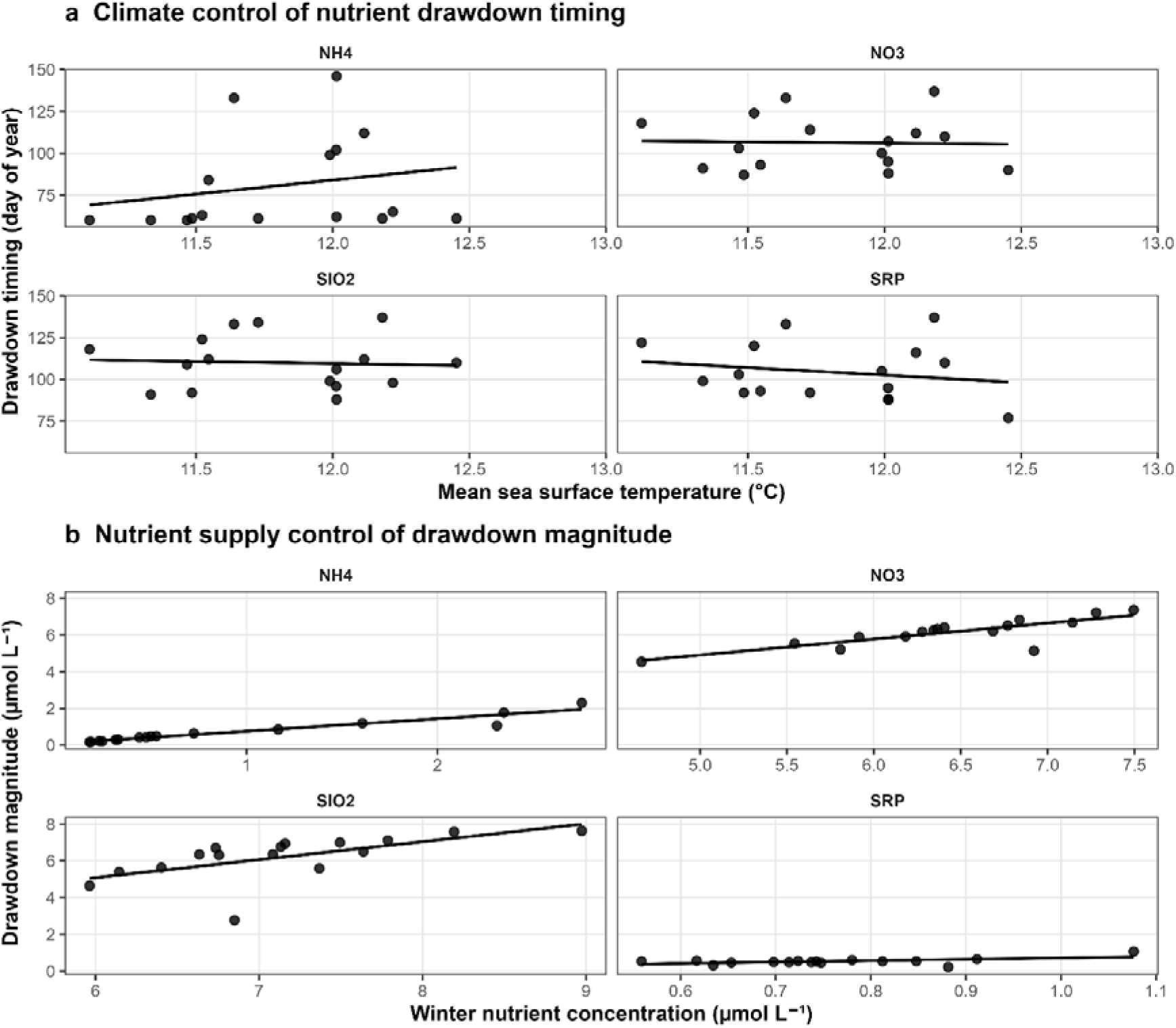
Timing and magnitude of seasonal nutrient drawdown. **a,** Relationships between mean sea surface temperature and the timing of seasonal nutrient drawdown (day of year) for nitrate (NO₃), ammonium (NH₄), phosphate (SRP), and silicate (SiO₂). Across nutrients, drawdown timing exhibited substantial interannual variability but showed only weak associations with sea surface temperature, suggesting that near-surface thermal conditions alone do not strongly constrain nutrient utilisation phenology. **b,** Relationships between winter nutrient concentration and subsequent drawdown magnitude show that seasonal nutrient utilisation is strongly constrained by pre-bloom nutrient inventories.

We tested how bloom intensity changed in response to differences in the magnitude of seasonal nutrient drawdown for nitrate (NO₃), silicate (SiO₂), and phosphate (SRP) (Figure 3; Supplementary Table 3). Bloom intensity showed weak and non-significant relationships with nitrate drawdown magnitude (R² = 0.001, p = 0.889), indicating that the extent of nitrate depletion alone does not explain interannual variability in bloom expression once drawdown has occurred. Bloom intensity increased with silicate drawdown magnitude (Figure 3b). Moderate positive relationships were detected between silicate drawdown and bloom intensity (R² = 0.24, p = 0.054), representing the strongest nutrient–bloom relationship observed across the three nutrients examined. Phosphate drawdown exhibited an intermediate pattern, with a positive association with bloom intensity that was weaker than that observed for silicate but stronger than for nitrate (R² = 0.104; p = 0.222) (Figure 3c). These results suggest that while nitrate drawdown is a prerequisite for bloom development, bloom intensity is more strongly influenced by silicate availability, with phosphate potentially playing a secondary role. Generalised additive models were fitted with restricted smoothers (k = 4), identifying weak nonlinear structure in the silicate–bloom relationship. However, these provided minimal improvement over linear models (ΔAIC = 0.33; Supplementary Table 3). GAM analyses similarly failed to improve model performance for nitrate drawdown, confirming the absence of hidden nonlinear structure in that relationship. The results, therefore, support the use of linear models to characterise first-order relationships between nutrient drawdown and changes in bloom expression.

**Figure 3.**
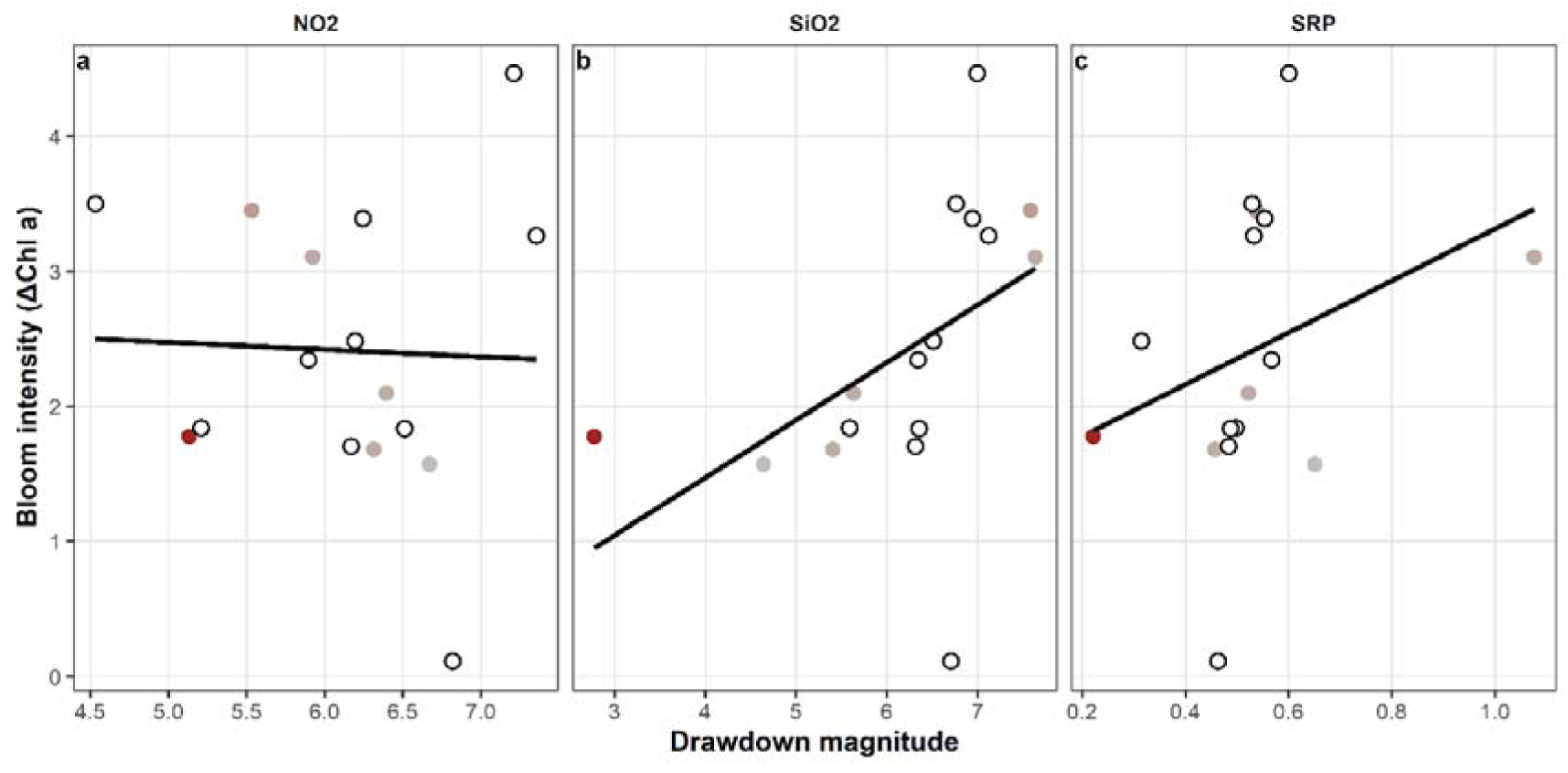
Bloom intensity as a function of the magnitude of seasonal drawdown for **a** nitrate (NO₃), **b** silicate (SiO₂), and **c** phosphate (SRP). The relationship between bloom intensity and drawdown magnitude differs among nutrients, with the strongest association observed for silicate. Point colour indicates diatom abundance (grey to red), and open circles denote observations where diatom abundance was unavailable.

Variations in bloom intensity for a given level of nutrient drawdown were associated with differences in phytoplankton community structure, as reflected by interannual variation in diatom abundance (Figure 3). Under silicate drawdown, years characterised by higher peak diatom abundance generally exhibited greater bloom intensity for a given drawdown magnitude, consistent with the obligate silicate requirement of diatoms and their contribution to the spring bloom. In contrast, nitrate drawdown showed little systematic relationship with either bloom intensity or diatom abundance. Phosphate drawdown exhibited an intermediate response, with bloom intensity increasing with drawdown magnitude and showing modest structuring by diatom abundance. Together, these patterns indicate that bloom expression is influenced not only by the extent of nutrient utilisation but also by phytoplankton community composition. While nutrient drawdown defines the potential for bloom development, realised bloom intensity depends on nutrient-specific physiological requirements and ecosystem structure rather than nutrient depletion alone.

Relationships between climatic variables and maximal diatom abundance were generally weak (Supplementary Figure 5). Linear relationships between mean sea surface temperature, cumulative heatwave intensity, and the winter NAO index explained little of the interannual variation in maximum diatom abundance (R² < 0.25 in all cases; Supplementary Table 3). In contrast to the relationships observed between nutrient drawdown, diatom abundance, and bloom intensity (Figure 3), climate metrics showed limited explanatory power for variation in bloom-forming biomass. The explanatory power of climate metrics, therefore, declined progressively from nutrient-system responses to biological responses. Climate explained up to 57% of the variance in nutrient drawdown timing and 43% of the variance in winter nutrient inventories, whereas relationships weakened for seasonal nutrient drawdown (31%) and bloom intensity (8%).

## Discussion

Within the western Irish Sea gyre, relationships between climate variability, nutrient cycling, and phytoplankton bloom dynamics weaken across successive physical, chemical and biological processes. Whilst we cannot improve the prediction of spring bloom intensity, our study provides a mechanistic explanation for why climate-bloom relationships remain weak despite strong physical and biogeochemical controls (Behrenfeld and Boss 2014, Payne et al. 2016, Planque 2016, Friedland et al. 2024). Climate variability influences key physical and biogeochemical processes, but its effects are progressively filtered through nutrient inventories, biogeochemical processes, and phytoplankton community structure (Behrenfeld and Boss 2014, Planque 2016). Consequently, weak relationships between climate variability and bloom intensity are an expected property of the system, rather than evidence that climate forcing is unimportant (Payne et al. 2016, Friedland et al. 2024).

We observed a separation between the controls on timing and magnitude of seasonal nutrient drawdown. Interannual variability in the onset of drawdown showed only weak associations with climate metrics, indicating that climate forcing exerts only a modest influence on the phenology of nutrient drawdown during a phytoplankton bloom (Gowen et al. 1995, Li et al. 2000, Barrera-Alba et al. 2019). By contrast, the magnitude of seasonal nutrient drawdown was strongly constrained by winter nutrient inventories, with winter concentrations explaining a large fraction of variability in total nutrient consumption. It is, therefore, the pre-bloom nutrient inventory that primarily determines nutrient consumption (Sharples et al. 2006, Jardine et al. 2022). By separating these timing and inventory controls our study helps to resolve long-standing ambiguities in bloom–nutrient relationships reported in shelf-sea systems (Cloern and Jassby 2008, Planque 2016). Phenological shifts in bloom onset or nutrient drawdown do not necessarily translate into larger blooms or enhanced nutrient consumption, because the potential for biological production is limited by winter nutrient availability (Sharples et al. 2006). As a result, years characterised by earlier or later drawdown timing can exhibit similar drawdown magnitudes, and conversely, large differences in drawdown magnitude can arise without pronounced shifts in timing (Planque 2016). This helps to explain why interannual shifts in bloom timing often show little correspondence with bloom magnitude in observational records (Friedland et al. 2024).

Our results suggest that the processes mediating seasonal access to nutrients can be partially decoupled from those determining the size of the available nutrient pool. This is consistent with previous studies that indicate partial decoupling between water column nutrient availability and nutrient supply to the phytoplankton (Sharples et al. 2006, Planque 2016). Winter mixing and the resulting nutrient inventories set an upper limit on phytoplankton bloom production, while climate-driven variability primarily affects the timing of the bloom (Jardine et al. 2022). Predictability is, therefore, strongest in the relationship between winter nutrient supply and seasonal drawdown and weakens as nutrient use is translated into phytoplankton biomass (Behrenfeld and Boss 2014, Payne et al. 2016). Even where seasonal nutrient drawdown is substantial, nitrate use alone provides little explanatory power for interannual variability in bloom intensity. Nitrate depletion is a necessary condition for spring bloom development in shelf-sea systems (Sverdrup 1953, Cullen 1991). However, it is not sufficient to accurately determine the magnitude of the phytoplankton bloom (Behrenfeld 2010). Once nitrate drawdown has occurred, transformation of the nutrient into biomass is constrained by factors that include nutrient stoichiometry and the functional requirements of different phytoplankton groups (Droop 1973, Arrigo 2005). The weak relationship between nitrate drawdown magnitude and bloom intensity indicates that bulk nitrogen consumption does not uniquely constrain biomass accumulation. Instead, the translation of nutrient use into phytoplankton biomass is likely mediated by nutrient stoichiometry and the functional requirements of different phytoplankton groups, including the availability of silicate for bloom-forming diatoms (Egge and Aksnes 1992, Arrigo 2005). Both linear and generalised additive models converged on this result, indicating that the weak nitrate-bloom relationship reflects an ecological property of the western Irish Sea ecosystem rather than a modelling artefact.

Silicate exhibited the strongest nutrient–bloom association observed in this study, although the relationship remained moderate and non-deterministic. Greater silicate drawdown was generally associated with higher bloom intensity, consistent with the obligate silicate requirement of bloom-forming diatoms (Egge and Aksnes 1992). Explanatory power was, however, limited and weakened at higher drawdown. This pattern indicates that silicate does not drive production. Instead, it acts as a functional constraint on the efficiency of nutrient use by phytoplankton (Droop 1973, Behrenfeld 2010). Thus, silicate availability shapes the phytoplankton bloom without uniquely determining its magnitude, reinforcing the view that bloom intensity is governed by interactions between nutrient stoichiometry and phytoplankton assemblage structure rather than control by a single resource (Arrigo 2005).

Variation in bloom intensity for a given level of nutrient drawdown highlights the role of biological structure as a key mediator of phytoplankton bloom dynamics (Behrenfeld et al. 2005, Behrenfeld 2010). In our study, we observed years with higher peak diatom abundance generally exhibited greater bloom intensity for a given silicate drawdown magnitude, consistent with the obligate silicate requirement of diatoms (Egge and Aksnes 1992), whereas this contextual structuring was weak or absent for nitrate drawdown (Behrenfeld 2010). Phosphate drawdown showed an intermediate response, with bloom intensity increasing with drawdown magnitude but only limited modulation by diatom abundance (Tilman 1977). Together, these observations indicate that nutrient drawdown establishes the potential for bloom development, while realised bloom intensity is shaped downstream by biological factors, including functional traits and food-web interactions, rather than determined solely by the extent of nutrient use (Irigoien et al. 2005, Behrenfeld 2010).

The relationships between climatic variables and maximum diatom abundance revealed consistently weak and variable associations, with low explanatory power across all cases. This indicates that, at interannual scales, phytoplankton community structure, as represented by diatom abundance, is not tightly predicted by climate forcing alone (Edwards and Richardson 2004). Instead, diatom abundance and associated bloom characteristics are emergent properties of the interactions of multiple processes operating downstream of climate variability, including nutrient inventories, stoichiometric constraints, and food-web interactions (Behrenfeld and Boss 2014). The absence of strong or consistent climate– diatom coupling does not imply a lack of physical influence. Instead, this likely reflects the indirect manner in which climate signals propagate through an ecosystem (Planque 2016). Thus, biological structure arises through emergent ecosystem dynamics rather than direct climatic controls (Behrenfeld and Boss 2014, Payne et al. 2016).

Our results suggest that direct prediction of phytoplankton bloom dynamics from climatic metrics alone is likely to remain challenging in shelf-sea ecosystems. Climate variability primarily influences bloom development by altering the timing and processes that regulate nutrient access, whereas bloom intensity reflects nutrient constraints and phytoplankton community structure (Behrenfeld 2010). Consequently, improved understanding of climatic forcing alone is unlikely to yield proportional gains in predictive skill for phytoplankton blooms at interannual timescales (Payne et al. 2016). This has important implications for ecosystem modelling, as even well resolved physical and biogeochemical drivers cannot fully account for the variability observed in biological responses of major phytoplankton groups (Behrenfeld and Boss 2014). From a management perspective, these findings caution against expectations of direct or predictable biological responses to climate change in shelf seas, as climatic signals are progressively modified by interacting physical, chemical and biological processes (Lotze et al. 2006, Planque 2016).

Whilst the dissolved nutrient and sea temperature records at the mooring extend back to 1997, our analysis was constrained by the availability of a consistent chlorophyll time series from 2006 onwards. Extension of biological observations over longer timescales may provide additional insight into lower-frequency ecosystem responses to climate variability and help resolve whether the attenuation processes identified here remain consistent across longer climatic cycles. Furthermore, drawdown magnitude is calculated relative to winter nutrient concentrations, meaning that some degree of statistical dependence between these variables is inherent to metric construction. The strength and consistency of the relationships, however, suggest that this is a genuine ecological response and not solely driven by mathematical dependence.

Overall, our results support a hierarchical model of climate-signal propagation within shelf-sea ecosystems. Climate variability influences nutrient availability and the timing of seasonal drawdown, but these effects become progressively attenuated as they pass through biogeochemical processes, phytoplankton community structure, and trophic interactions. Consequently, weak climate-bloom relationships should be viewed not as a failure to detect climatic forcing, but as an emergent property of ecosystem organisation. Whilst focused on the western Irish Sea gyre, the processes identified here are unlikely to be unique to this system. Seasonally stratified shelf seas worldwide share common characteristics, including finite winter nutrient inventories, pronounced seasonal transitions in physical structure, and strong biological mediation of nutrient use (Sharples et al. 2006, Bauer et al. 2013). The western Irish Sea therefore functions as a model system in which these interactions can be resolved clearly, suggesting that attenuation of climatic signals during their propagation through successive ecosystem processes may be a general feature of temperate shelf-sea ecosystems (Planque 2016, Friedland et al. 2024). Weak and variable climate–bloom relationships should therefore be expected across many shelf-sea systems (Behrenfeld and Boss 2014, Payne et al. 2016). Improving our predictions of phytoplankton bloom dynamics may, therefore, depend less on improved characterisation of climatic forcing and more on understanding the ecological and biogeochemical processes that mediate nutrient utilisation and biomass accumulation.

## Acknowledgements

We thank the masters and crew of RV Corystes for their support in maintaining the western Irish Sea time-series programme. Long-term observational records are built upon the dedication of multiple generations of scientists and technical staff, and we gratefully acknowledge the contributions of current and former AFBI personnel, including Brian Stewart, Richard Gowen, Chris Gibson, Raymond Gilmore, Patrick Quinn, and many others whose efforts have sustained this programme over more than three decades. This work is funded by the Northern Ireland Department of Agriculture, the Environment and Rural Affairs through AFBI’s mission to provide Science to Underpin an Ecosystem Approach to Sustainable Management of Irish Sea Fisheries.

