## Supplementary Information for "Winter nutrients constrain spring drawdown, but only weakly predict bloom intensity in the western Irish Sea"

### Supplementary Methods: Validation of Reconstructed Nutrient Time Series

To evaluate the robustness of the post‑2023 reconstructed nutrient time series, nutrient concentrations obtained from the fixed‑depth McLane® Remote Access Sampler (RAS‑100) were compared with contemporaneous ship‑based CTD/rosette observations collected during periods of temporal overlap. Ship‑based CTD casts were conducted approximately every six weeks during routine servicing of the Station 38A mooring. Nutrient concentrations derived from CTD rosette sampling represent depth‑averaged values over the upper 0–40 m water column, whereas McLane observations represent fixed‑depth measurements collected at approximately 20 m depth. To assess the consistency of these measurement approaches and quantify any systematic offsets, paired observations were extracted where CTD sampling occurred within ±1 day of McLane sampling.

For each nutrient (NO₃⁻, NH₄⁺, SRP, SiO₂, and NO₂⁻), ordinary least‑squares linear regression models were fitted:

$$\text{CTD}_{i}=\alpha+\beta\text{ }\text{McLane}_{i}+\varepsilon_{i}$$

where $\mathrm{CTD}_{i}$ represents the depth‑integrated CTD concentration and $\mathrm{McLane}_{i}$ the corresponding fixed‑depth moored observation. Regression coefficients, coefficients of determination (R²), root mean square error (RMSE), and residual diagnostics were examined for each nutrient to evaluate: (i) the strength of correspondence between sampling approaches; (ii) the presence of systematic offsets or proportional bias, and; (iii) the suitability of the regression relationships for bias correction of CTD observations in the post‑2023 period. The residuals from each regression were visually assessed and showed no evidence of systematic non‑linearity or heteroscedasticity; these diagnostics are therefore not shown.

The results of the regressions analysis are summarised in supplementary figure 4 and supplementary table 3. A strong linear relationship (R² = 0.87) was observed for nitrate, indicating that both sampling approaches resolve a common underlying seasonal signal dominated by winter accumulation and spring drawdown. Moderate linear relationships were detected for both silicate (R² = 0.41) and soluble reactive phosphorus (R² = 0.45), characterised by increased scatter and lower coefficients of determination (R² = 0.41). These differences suggest tighter coupling of silicate and SRP concentrations to short‑term biological uptake, rapid recycling, and sub‑seasonal variability, which reduce pointwise correspondence between discrete bottle samples and time‑integrated moored observations. Reduced pointwise agreement for silicate and SRP does not imply the absence of a coherent seasonal cycle. The reconstructed climatology represents the mean seasonal trajectory of nutrient availability rather than instantaneous concentrations. Consequently, it is used conservatively to define seasonal structure, phenology, and relative changes through time, rather than absolute concentration values. Bias‑corrected CTD values were, therefore, only applied where regression relationships were well constrained, while climatological estimates were retained for unsampled periods and for nutrients where correction would risk imposing artificial structure.

**Supplementary Figure 4** Comparison of fixed‑depth McLane® sampler and depth‑integrated CTD bottle nutrient concentrations during periods of temporal overlap at Station 38A. Points represent paired observations (±1 day), with solid lines indicating ordinary least‑squares regression fits and dashed lines denoting 1:1 correspondence. Regression strength varies among nutrients, with nitrate showing strong linear agreement and other nutrients exhibiting greater scatter. Reduced pointwise correspondence for silicate and soluble reactive phosphorus reflects increased short‑term biological variability and sampling‑scale differences, rather than systematic measurement bias, and does not preclude the use of climatological seasonal structure for phenological analyses.

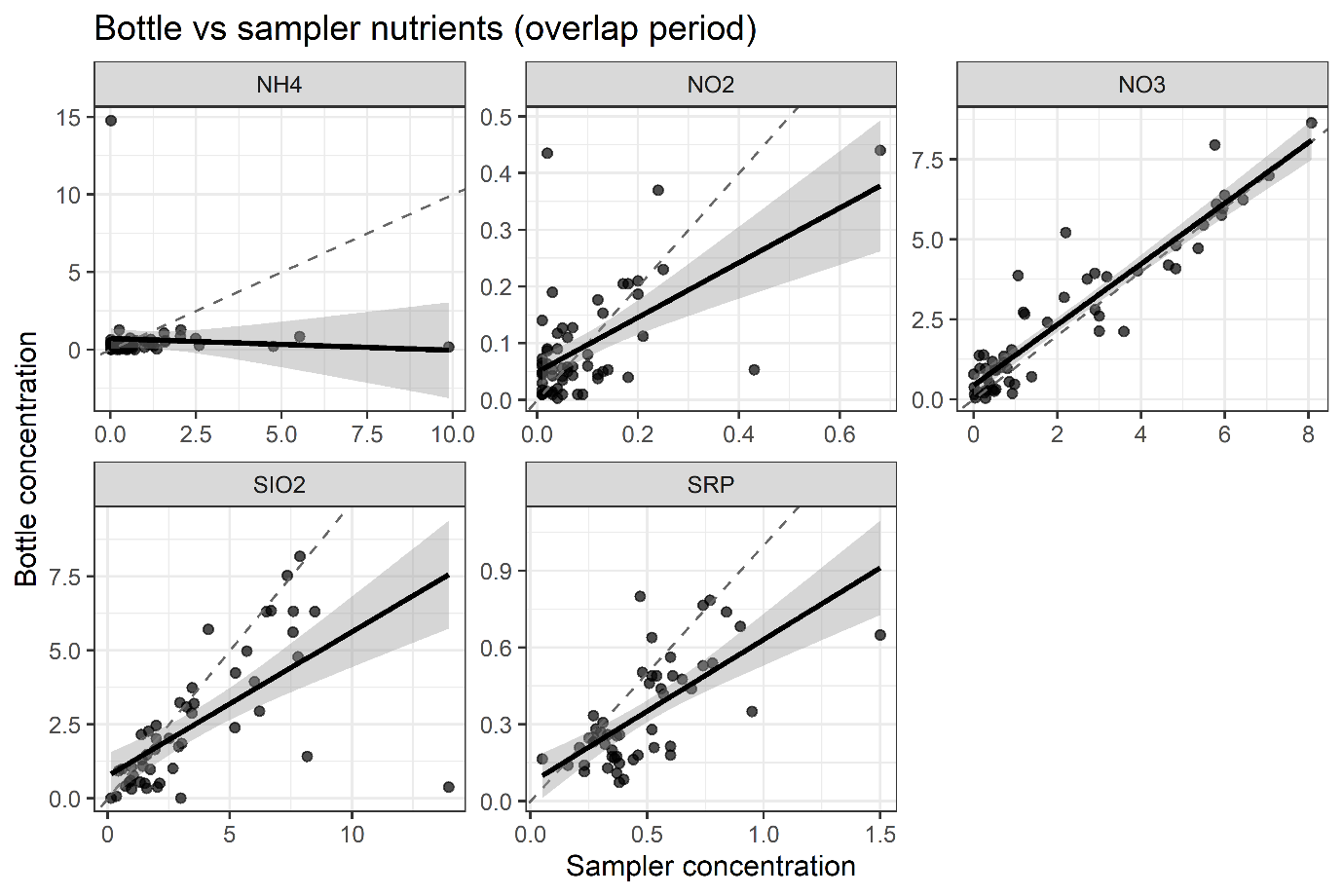

**Supplementary Table 1.** Linear regression diagnostics comparing fixed‑depth McLane® sampler nutrient concentrations with depth‑integrated CTD bottle measurements during periods of temporal overlap. Metrics summarise pointwise correspondence between sampling approaches and were used to assess the suitability of individual nutrients for bias correction versus conservative representation using a seasonal climatology.

| **Nutrient** | **n** | **Slope** | **Intercept** | **R²** | **RMSE** | **Correspondence strength** | **Used for bias correction** |
| --- | --- | --- | --- | --- | --- | --- | --- |
| NO₃⁻ | 56 | 0.95 | 0.43 | 0.87 | 0.81 | **Strong** | **Yes** |
| SiO₂ | 50 | 0.49 | 0.76 | 0.41 | 1.67 | Moderate | No |
| SRP | 51 | 0.56 | 0.07 | 0.45 | 0.15 | Moderate | No |
| NO₂⁻ | 57 | 0.48 | 0.05 | 0.32 | 0.08 | Weak–moderate | No |
| NH₄⁺ | 53 | −0.08 | 0.72 | <0.01 | 1.98 | Weak | No |

**Supplementary Table 2.** Comparison of linear models (LMs) and generalized additive models (GAMs) describing relationships between phytoplankton bloom intensity and seasonal nutrient drawdown metrics in the western Irish Sea. For each response-predictor combination, model performance is summarised by coefficient of determination (R²) and p-value for linear models, and effective degrees of freedom (EDF) and deviance explained for GAMs. Model fit was compared using Akaike's Information Criterion (AIC), with positive ΔAIC values indicating improved support for the GAM relative to the corresponding linear model.

| Response variable | Predictor variable | n | LM R² | LM p-value | LM AIC | | GAM EDF | GAM deviance explained (%) | GAM AIC | ΔAIC  (LM − GAM) |
| --- | --- | --- | --- | --- | --- | --- | --- | --- | --- | --- |
| Bloom intensity | Silicate drawdown magnitude | 16 | 0.24 | 0.054 | 47.83 | 1.67 | | 34.5 | 47.5 | 0.33 |
| Bloom intensity | Nitrate drawdown magnitude | 16 | 0.001 | 0.889 | 52.2 | 1.5 | | 9.1 | 52.35 | -0.15 |

### Supplementary Results

**Supplementary Figure 1 a,** Time series of winter nutrient concentrations and seasonal drawdown magnitude for nitrate (NO₃), ammonium (NH₄), phosphate (SRP), silicate (SiO₂), and nitrite (NO₂). Across nutrients, winter availability and drawdown magnitude exhibit coherent interannual variability, with nutrient‑specific differences in amplitude. **b,** Time series of physical climate variables used in the analysis, including mean sea surface temperature, cumulative heatwave intensity, cumulative cold‑spell intensity, and winter‑mean North Atlantic Oscillation (DJFM). These data illustrate the range of climate variability considered and provide context for the mechanistic analyses presented in the main text.

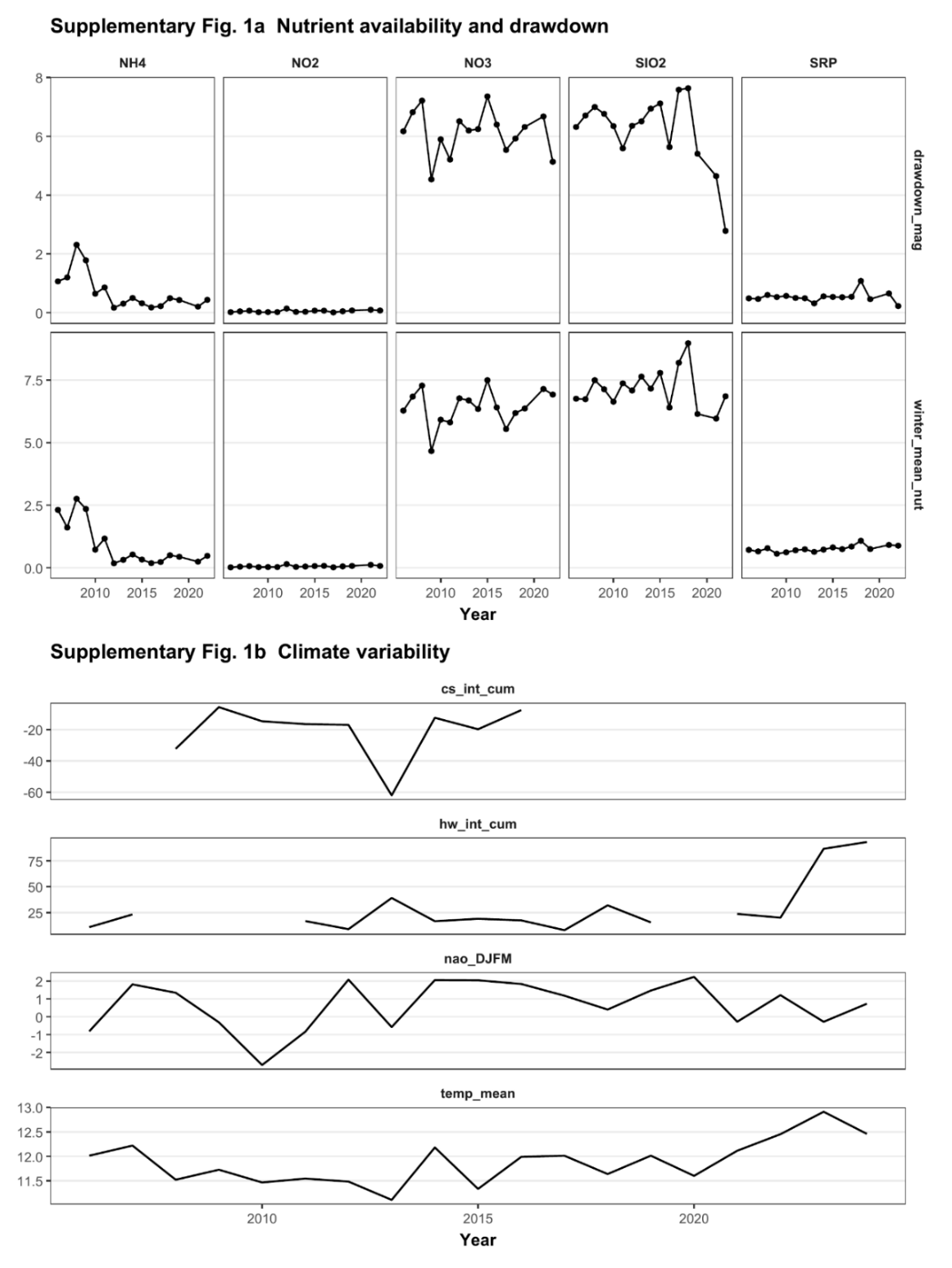

**Supplementary Figure 2.** Additional analyses of nutrient drawdown dynamics. **a,** Relationships between nutrient drawdown timing and multiple climate metrics, including mean sea surface temperature, cumulative heatwave intensity, and winter‑mean North Atlantic Oscillation (DJFM), shown separately for each nutrient. Independent x‑axis scaling highlights the differing physical ranges of climate variables while demonstrating consistent nutrient‑specific responses. **b,** Relationships between drawdown magnitude and spring minimum nutrient concentration for each nutrient. Greater drawdown magnitude is associated with lower spring minimum concentrations, indicating that seasonal depletion directly structures post‑bloom nutrient availability.

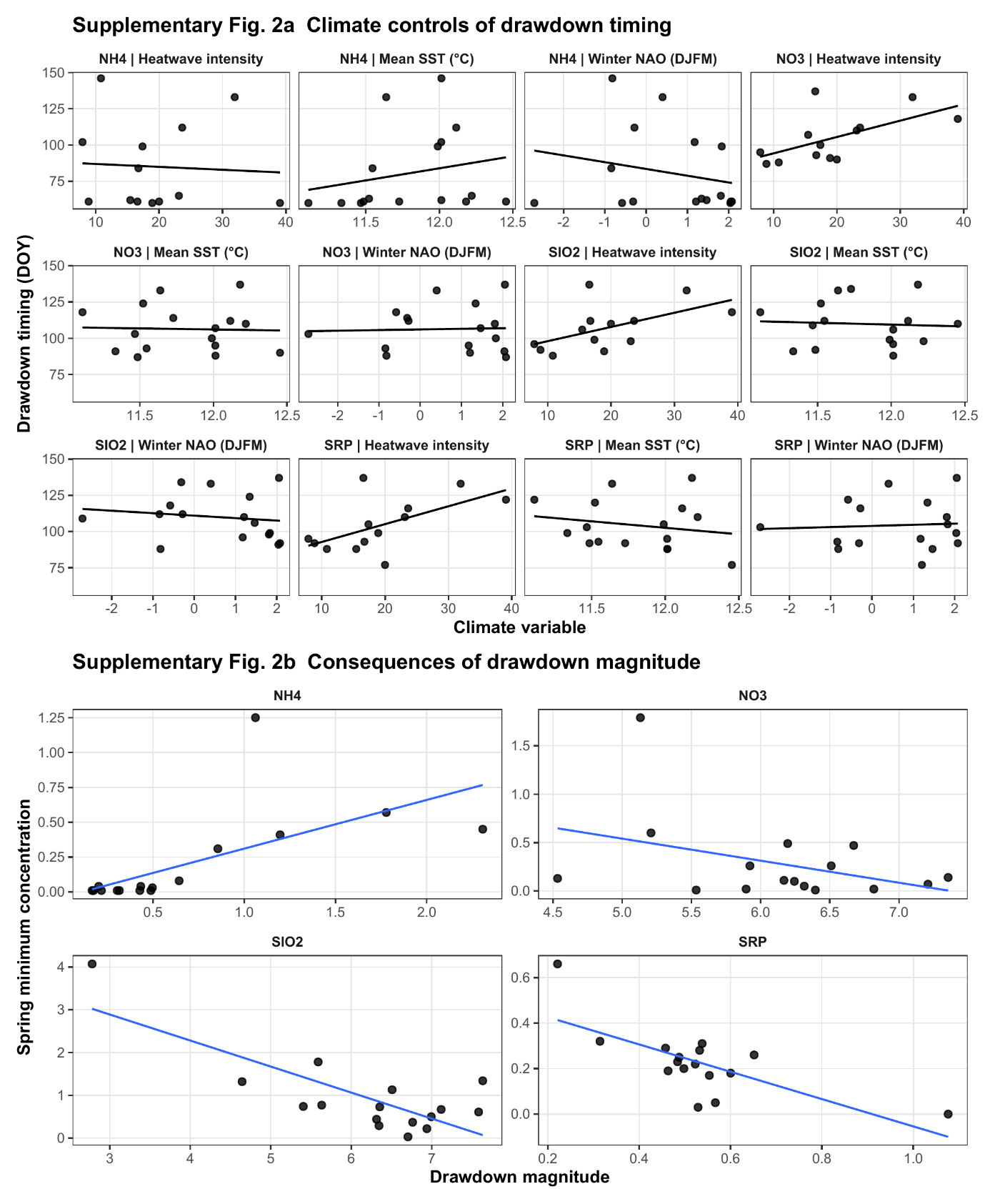

**Supplementary Figure 3** Relationships between **a** cumulative heatwave intensity, **b** mean sea surface temperature, and **c** winter‑mean North Atlantic Oscillation and maximum diatom abundance. Across metrics, associations are weak and variable, indicating that interannual variability in diatom biomass is not tightly prescribed by large‑scale climate forcing.

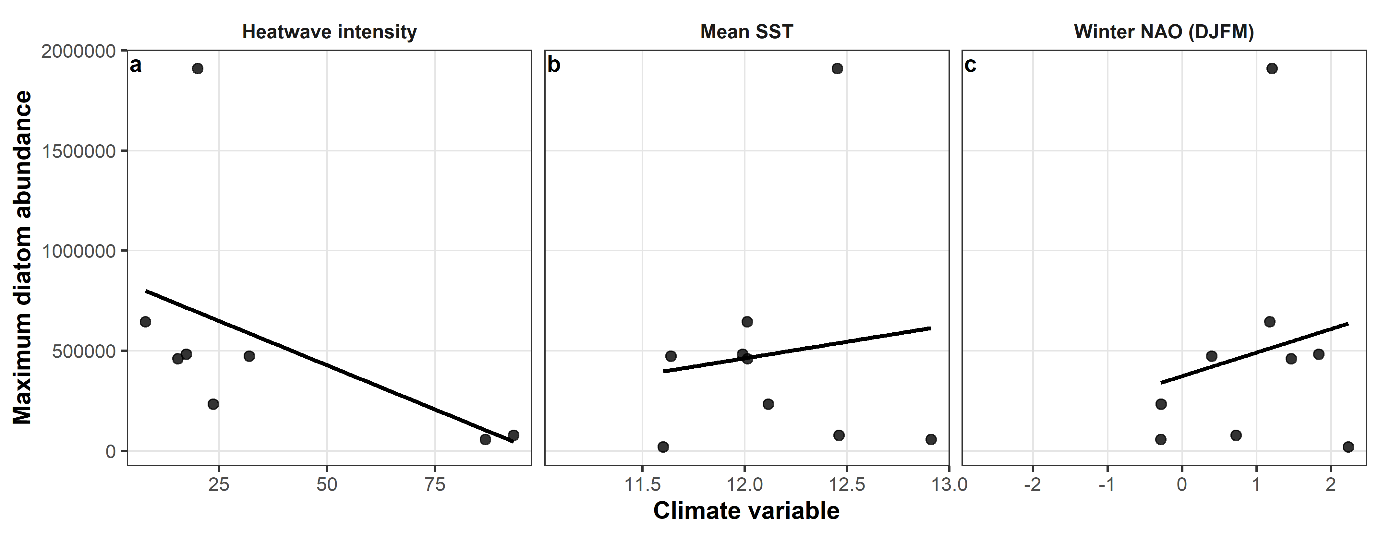

**Supplementary Table 3**. Linear regression results testing relationships between climate, nutrient dynamics, and phytoplankton bloom metrics in the western Irish Sea. Values shown are regression slope, intercept, coefficient of determination (R²), and significance level (*p*).

| Response | Predictor | Slope | Intercept | r^2^ | p |
| --- | --- | --- | --- | --- | --- |
| Ammonium Drawdown timing | Mean SST | 16.69 | -116.347 | 0.047 | 0.422 |
| Nitrate Drawdown timing | Mean SST | -1.485 | 123.9043 | 0.001 | 0.898 |
| Silicate Drawdown timing | Mean SST | -2.509 | 139.5459 | 0.004 | 0.827 |
| Phosphorus Drawdown timing | Mean SST | -8.94 | 209.8878 | 0.038 | 0.471 |
| Ammonium Drawdown magnitude | Winter nutrient concentration | 0.676 | 0.0872 | 0.912 | <0.0001 |
| Nitrate Drawdown magnitude | Winter nutrient concentration | 0.869 | 0.5586 | 0.671 | <0.0001 |
| Silicate^-^ Drawdown magnitude | Winter nutrient concentration | 0.977 | -0.7747 | 0.383 | 0.0106 |
| Phosphorus Drawdown magnitude | Winter nutrient concentration | 0.777 | -0.0586 | 0.311 | 0.0249 |
| Bloom intensity | NOx- Drawdown magnitude | -0.053 | 2.7394 | 0.001 | 0.889 |
| Bloom intensity | SiO2- Drawdown magnitude | 0.427 | -0.23613 | 0.24 | 0.0541 |
| Bloom intensity | SRP Drawdown magnitude | 1.92 | 1.393806 | 0.104 | 0.222 |
| Diatom*_max_* | MHW Cumulative Intensity | -8837.17 | 868444.6 | 0.25 | 0.207 |
| Diatom*_max_* | winter NAO | 117968 | 372301.8 | 0.032 | 0.645 |
| Diatom*_max_* | Mean SST | 165532.1 | -1525092 | 0.014 | 0.76 |
